# A data-driven method for reconstructing and modelling social interactions in moving animal groups

**DOI:** 10.1101/816777

**Authors:** R. Escobedo, V. Lecheval, V. Papaspyros, F. Bonnet, F. Mondada, C. Sire, G. Theraulaz

## Abstract

Group-living organisms that collectively migrate range from cells and bacteria to human crowds, and include swarms of insects, schools of fish and flocks of birds or ungulates. Unveiling the behavioural and cognitive mechanisms by which these groups coordinate their movements is a challenging task. These mechanisms take place at the individual scale and they can be described as a combination of pairwise interactions between individuals and interactions between these individuals and the physical obstacles in the environment. Thanks to the development of novel tracking techniques that provide large and accurate data sets, the main characteristics of individual and collective behavioural patterns can be quantified with an unprecedented level of precision. However, in a large number of works, social interactions are usually described by force map methods that only have a limited capacity of explanation and prediction, being rarely suitable for a direct implementation in a concise and explicit mathematical model. Here, we present a general method to extract the interactions between individuals that are involved in the coordination of collective movements in groups of organisms. We then apply this method to characterize social interactions in two species of shoaling fish, the rummy-nose tetra (*Hemigrammus rhodostomus*) and the zebrafish (*Danio rerio*), which both present a burst-and-coast motion. The detailed quantitative description of microscopic individual-level interactions thus provides predictive models of the emergent dynamics observed at the macroscopic group-level. This method can be applied to a wide range of biological and social systems.

## 1 Introduction

The identification and characterization of interactions between the constituent elements of a living system is a major challenge for understanding its dynamic and adaptive properties [1, 2]. In recent years, this issue is also at the heart of research conducted in the field of collective behaviour in animal groups and societies [3, 4]. The identification from field data of the interaction rules between individuals in species whose level of cognitive complexity can be quite high remains problematic. Indeed, the way in which individuals interact is strongly influenced and modulated by the physical characteristics of the environment in which the organisms live, such as temperature, humidity, brightness, or even by the presence of air currents (see for instance [5, 6] in social insects). The situation is quite different in the laboratory, where conditions can be precisely controlled. Moreover, new tracking techniques make it possible to record the behaviour of individuals alone or in groups along relatively long periods of time [7, 8, 9, 10, 11]. Using large sets of tracking data, one can then reconstruct and model the social interactions between two individuals of the same species and between them and the obstacles present in their environment [12, 13].

Explaining collective behaviour in groups of organisms consists in describing the mechanisms by which the behaviour of an individual is influenced by the behaviour of the other group members that are present in its neighbourhood [14, 15]. The output of the behaviour of an individual is the set of its successive positions during a given period of time (usually, at some discrete time steps). From these data, it is possible to draw the trajectories of all the individuals in a group and calculate their instantaneous velocity and acceleration, their distance and angle of incidence to obstacles (or the wall in an experimental tank), their heading, as well as relative quantities such as the distance between individuals and the angle of their relative position, and group quantities such as cohesion and heading polarization. These measures can reveal individual behavioural patterns such as the average velocity of motion or the frequency of heading changes close to obstacles, and also collective behavioural patterns such as the level of polarization. Thus, the analysis of collective behaviour consists in measuring behavioural changes at the individual scale that likely result from social interactions, and to associate these measures with the relative state of the individuals involved in these interactions [12]. The relative state of an individual *j* with respect to a focal individual *i* is determined by the distance *d*_*ij*_ between *i* and *j*, its relative velocity v_*ij*_, its relative position with respect to *i*, *ψ*_*ij*_, and its relative heading *φ*_*ij*_ (see Fig. 1). The behavioural changes are precisely given by the variations of an individual’s position and velocity, or, equivalently, by the position, speed and heading variations.

**Figure 1:**
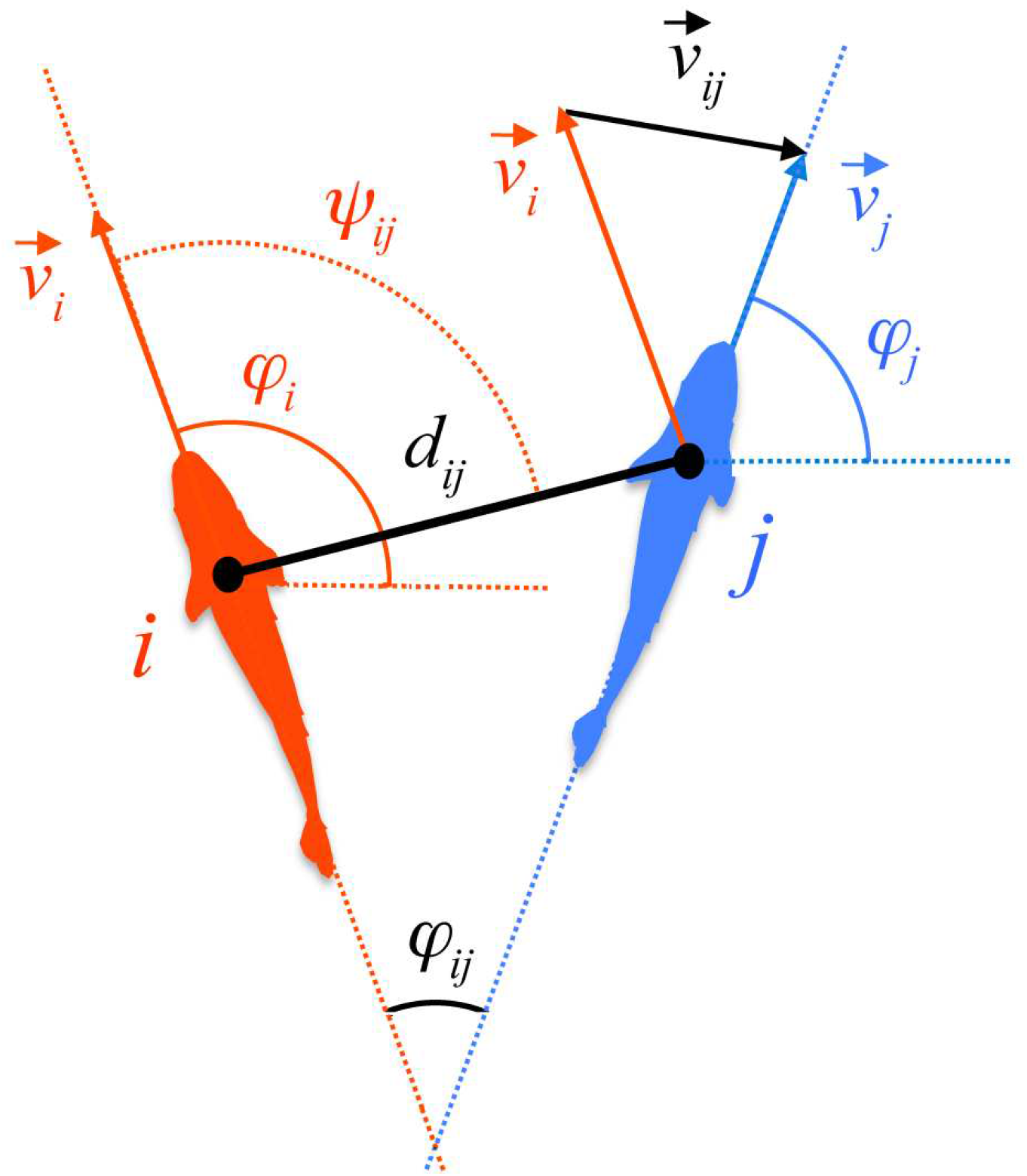
State variables of a focal fish *i* with respect to its neighbour *j*. Fish position is determined by the position of the centre of mass of the fish (black circles) in a orthonormal system of reference *Oxy*. *d*_*ij*_: distance between fish; 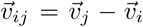: relative velocity of *j* with respect to *i*; *ψ*_*ij*_: angle with which *i* perceives *j*; *φ*_*ij*_ = *φ*_*j*_ − *φ*_*i*_: relative heading of *j* with respect to *i*. Angles are measured with respect to the horizontal axis of coordinates *Ox*; we use the convention that angles are positive in the counterclockwise direction.

In fish that have a burst-and-coast swimming mode, behavioural changes correspond to significant variations of the individual’s heading that occur exactly at the onset of the acceleration phase (*i.e.*, the bursts). These discrete behavioural decisions are called “kicks” [12, 16]. Other quantities such as the intensity of the acceleration in the direction perpendicular to the direction of motion, or simply the turning direction (right or left), can be used to detect behavioural changes. The task is thus to put in relation the heading variation of a focal fish *δφ*_*i*_ with its state variables, that is, to find a function *δφ*_*i*_(*d*, v, *ψ, φ*).

In this article, we first show that force maps, that are widely used to describe the effects of social interactions on the behaviour of individuals, have important limitations when it comes to model the interactions between individuals. These limitations result mostly from the limited number of variables that force maps can handle, the difficulty of identifying intermediate contributions to behavioural patterns, and the difficulty of distinguishing the effects of state variables from constitutive parameters. We then describe in detail a method to analyse behavioural data obtained from digitized individual trajectories. The method allows us (1) to quantify the social interactions between two individuals and describe how the intensities of these interactions vary as a function of the state variables of the individuals, and, (2) to reconstruct analytically the interaction functions and to derive an explicit and concise mathematical model reproducing the observed behaviours. Finally, we apply this method to the analysis of the social interactions in two species of fish that both have a burst-and-coast type of swimming and that are characterized by very different levels of coordination when swimming in groups. The reconstruction of interaction rules allows to understand the origin of the differences in the level of coordination, and to predict in which experimental conditions other behavioural differences can arise.

## 2 Use and limitations of force maps to infer social interactions

A first way to infer social interactions directly from experimental data consists in using the force-map technique [41]. This technique has been used for instance to estimate from experiments performed with two fish the effective turning and speeding forces experienced by an individual, once the relevant variables on which they may depend have been chosen [11, 17, 18]. This is a simple way to visualise the strength and direction of behavioural changes. However, force maps have strong limitations that can induce profound misunderstandings.

### 2.1 Visualisation of social interactions with force maps

Force maps are colour maps, that is, 3D representations of 2D functions of the form *f* (*x, y*), where the variation of the value of the function is represented by a colour gradient. Generating and interpreting force maps is easy and this explains their success for inferring interactions between moving groups of individuals.

A 2D function *f* (*x, y*) given by a data set is a sequence of *N* triplets (*x*^*n*^, *y*^*n*^, *f*^*n*^), where the index *n* denotes for example the instant of time *t*^*n*^, *n* = 1, …, *N*. To build a force map of this function, the (*x, y*)-space is discretised in *I* × *J* boxes of the form 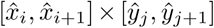, where the nodes 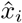 and 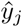 are given by

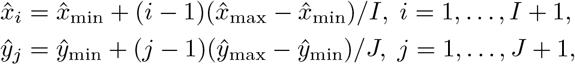

and the data (*x*^*n*^, *y*^*n*^, *f*^*n*^) are placed in a *ij*-box in such a way that 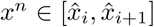 and 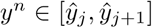. Then, the number of data *ϵ*_*ij*_ in the *ij*-box and the mean value *f*_*ij*_ of the values of *f*^*n*^ that fell in the *ij*-box are calculated. The value *f*_*ij*_ is then considered as the value of *f* (*x, y*) at the middle point of the *ij*-box, 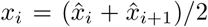, 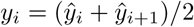. The resulting points (*x*_*i*_, *y*_*j*_, *f*_*ij*_) are then represented in a colour surface, after optional interpolation with, for instance, multilevel B-splines [19].

Figs. 2 and 3 show the force maps of the heading variation *δφ* of an individual fish as a function of different variables when the fish swims with a conspecific in a circular tank. For the case of *Hemigrammus rhodostomus*, we used a tank of radius *R* = 0.25 m, about 8.3 times the body length (BL) of the fish, and in the case of *Danio rerio*, a tank of radius *R*_Z_ = 0.29 m, about 6.4 BL. In both Figs. 2 and 3, panels A show the intensity of *δφ* as a function of *d*_*ij*_ and *ψ*_*ij*_, the distance between fish and the angle with which fish *i* perceives fish *j*, respectively, and panels B show *δφ* as a function of *d*_*ij*_ and *φ*_*ij*_, the heading difference between fish.

**Figure 2:**
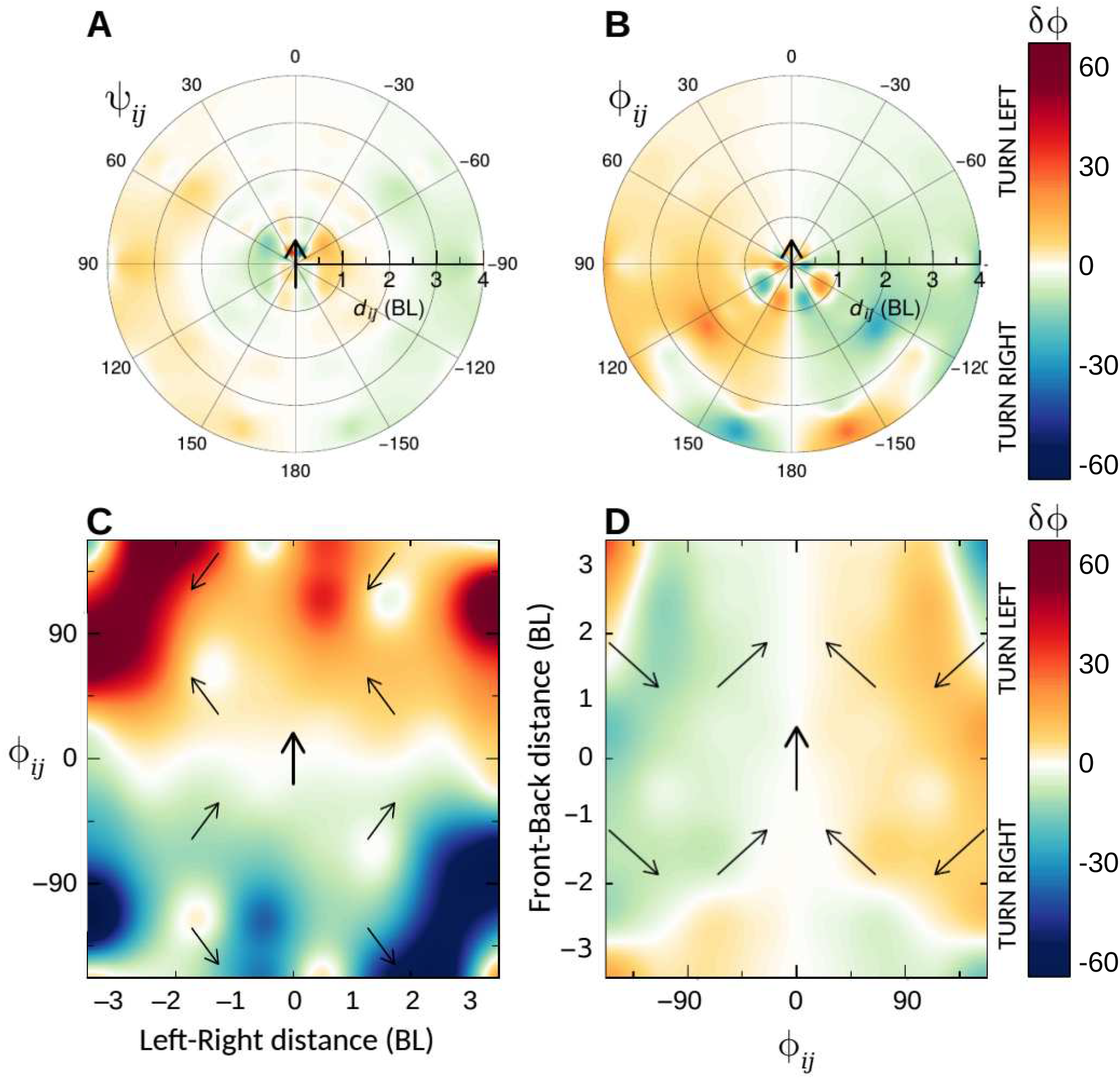
Force maps of *H. rhodostomus* swimming in a circular arena of radius 0.25 m. Heading change *δφ* of a focal fish, located at the origin and pointing north in the four panels, as a function of (A) the distance to its neighbour *d*_*ij*_ and its relative heading *φ*_*ij*_, (B) the distance *d*_*ij*_ and the angle of perception of its neighbour *ψ*_*ij*_, (C) the left-right distance 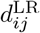 and *φ*_*ij*_, and (D) the relative heading *φ*_*ij*_ and the front-back distance 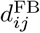. The colour scale shown in the right represents the average value of heading change *δφ*. Distances are measured in body lengths (BL), angles in degrees.

**Figure 3:**
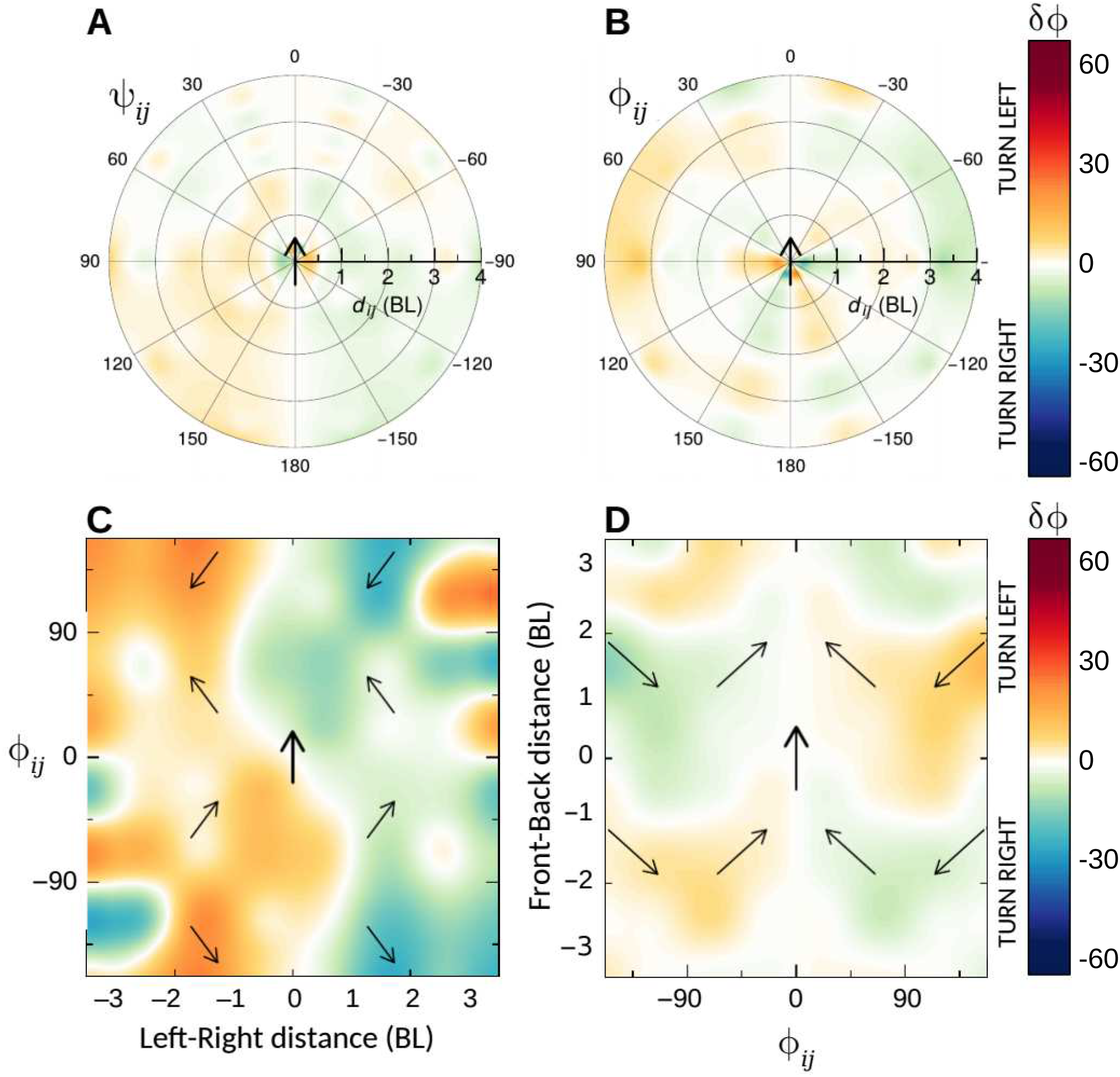
Force maps of *D. rerio* swimming in a circular arena of radius 0.29 m. Heading change *δφ* of a focal fish, located at the origin and pointing north in the four panels, as a function of (A) the distance to its neighbour *d*_*ij*_ and its relative heading *φ*_*ij*_, (B) the distance *d*_*ij*_ and the angle of perception of its neighbour *ψ*_*ij*_, (C) the left-right distance 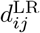 and *φ*_*ij*_, and (D) the relative heading *φ*_*ij*_ and the front-back distance 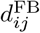. The colour scale shown in the right represents the average value of heading change *δφ*. Distances are measured in body lengths (BL), angles in degrees.

These force maps provide some information about the individual behaviour of fish. In *H. rhodostomus*, Fig. 2A shows that the focal fish tends to turn towards its neighbour, to the left (resp. right) when the neighbour is on the left (resp. right), except when the neighbour is very close. In that case, the behaviour is quite complex, the fish performs small angular changes of amplitude ≈ 30–60°, probably due to collision avoidance manoeuvres. When the neighbour is further from the focal fish, the latter maintains its heading (white circular region at *d*_*ij*_ ≈ 1–2 BL). From this force map, one could conclude that a fish is attracted by its neighbour when it is beyond a distance of about 2 BL, and repulsed when they are too close from each other (*d*_*ij*_ *<* 1 BL). The force map in Fig. 2B shows that the focal fish turns left (resp. right) when the relative heading of the neighbour is shifted to the left (resp. right). The larger the heading difference, the stronger the turn: the colour intensity increases as |*φ*_*ij*_| grows from 0 to 120°. When fish swim in more or less opposite directions, the intensity of the heading change is small. This force map reveals that the focal fish tends to align with its neighbour. In *D. rerio*, Fig. 3A shows that the focal fish turns towards its neighbour when their are close to each other (*d*_*ij*_ ≈ 1–2 BL) or when its neighbour is located behind the focal fish (|*ψ*_*ij*_| *>* 90°, whatever the distance). When the neighbour is at *d*_*ij*_ ≈ 2–3 BL in front of the focal fish (|*ψ*_*ij*_ *<* 60°), the latter turns away from its neighbour, as well as when the neighbour is very close to it (*d*_*ij*_ *<* 0.5 BL). The reaction to neighbour’s heading is less intense than in *H. rhodostomus*, as shown by the wide white regions in Fig. 3B. When fish are beyond 3 BL from each other, the focal fish turns to adopt the same orientation than its neighbour. At short distances, the behaviour is more complex and with changes of smaller size than those observed in *H. rhodostomus*.

The visualisation of the data by means of force maps therefore suggests the presence of two distinct types of interaction: an attraction interaction, where the fish turns towards its neighbour to get closer to it, and an alignment interaction, where the fish turns to adopt the same heading than its neighbour. However, nothing can be said from these colour maps about what happens when both contributions to heading variation have different sign: would a fish turn right or left when its neighbour is on its left side? Attraction alone would induce the focal fish to turn left. However, if the relative heading of the neighbour is turned to the right, alignment alone would induce the focal fish to turn right. This difficulty comes from the fact that a function (*δφ*) that depends on three variables (*d*_*ij*_, *ψ*_*ij*_, and *φ*_*ij*_) cannot be represented in 3D. To overcome this limitation, some authors use a kind of force map where the relative position of a fish with respect to a focal fish is decomposed in the left-right (LR) and front-back (FB) distances [13]. In Fig. 2A, (*d*_*ij*_, *ψ*_*ij*_) are the polar coordinates of fish *j* in the system of reference centred on fish *i* pointing north. This is a continuous system of reference where all the relative positions can be represented. Instead, LR and FB distances are projections of the relative position on the *d*_*ij*_-axis, where all the points of Fig. 2A that are in the left semicircle of radius 1 BL are averaged in a single point where 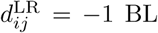 (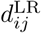 is the LR distance of fish *j* with respect to fish *i*), because these points are at 1 BL to the left of the focal fish. Then, the third variable *φ*_*ij*_ is used to expand this averaged point in a vertical line with different values of *φ*_*ij*_, in a system of reference with coordinates (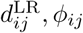), giving rise to Fig. 2C. Similarly, upper semicircles of Fig. 2A are averaged in the 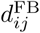-axis in Fig. 2D.

Force maps in panels C and D of Figs. 2 and 3 provide additional information about individual fish behaviour. In *H. rhodostomus*, Fig. 3C shows that turning direction is homogeneously distributed in the upper and lower half-planes, meaning that the focal fish turns to adopt the heading of its neighbour almost independently of the LR distance separating them, although the intensity of the turn is larger when the neighbour is far from the focal fish and perpendicular to it (regions of highly intense colour at 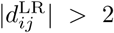 and *φ*_*ij*_ ≈ ±90°). In the white horizontal region, the focal fish maintains its heading when it is aligned with its neighbour (|*φ*_*ij*_ *<* 10°|), whatever the horizontal distance between them. Fig. 2D exhibits two large regions homogeneous in colour, showing that the focal fish turns almost always to adopt the heading direction of its neighbour, except when this one is far behind it (small regions of the opposite colour for 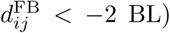). In *D. rerio*, the colour of each vertical half-plane of Fig. 3C is almost uniform, except for some regions close to the focal fish (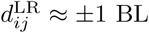) at *φ*_*ij*_ ≈ ±90°, and some distant regions located at 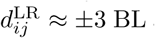 and *φ*_*ij*_ ≈ ±135°). This means that the focal fish turns almost always towards its neighbour, almost independently from their relative heading, except when they are close and perpendicular to each other, and when they are very far and almost anti-aligned. In Fig. 3D, one can see that the fish turns to adopt the same direction as that of its neighbour when this one is close and in front of it (large green and orange homogeneous regions in the centre of the figure). But the fish tends to turn towards the opposite direction when its neighbour is far ahead (small regions where 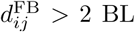) or behind it and not very close to it (regions where 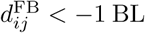). This is a different and more complex behaviour than the one observed in *H. rhodostomus*, where heading changes depend on *φ*_*ij*_ but not on 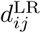. In *D. rerio*, it is the opposite, and the dependence on the FB distance is more complex than in *H. rhodostomus*. However, neither 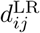, nor 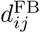, separately, can determine the distance at which the neighbour is (*e.g.*, a fish *j* located at 1 BL to the left can be at 0.5 or 2 BL to the front). Therefore, this kind of representation hides the effect of the absolute intensity of the interactions as a function of the distance between fish, and moreover does not allow to disentangle the contribution of intermediate effects such as attraction and alignment.

### 2.2 Limitations of force maps

The above descriptions show that the use of force maps to characterize interactions between individuals raises several problems, especially when more than two state variables must be taken into account. Other issues are not exclusive of force maps. For example, data can be scarce; not only because they are difficult to collect, but also because the phenomenon under observation rarely produce data of a given kind. This is what happens for example with repulsive interactions only: as individuals repel each other, there are very few instances in which the distance between them is small, so that repulsive interactions are very difficult to describe at short ranges. More importantly, data are rarely homogeneously distributed, so that different regions of the map can result from the average of a very disparate number of data, meaning that similar colour intensities do not have the same relevance. For example, Fig. 4 shows that the relative position of the neighbour of fish *i* is not homogeneously distributed on the (*d, ψ*)-plane, and is differently distributed in both panels, so that information from, *e.g.*, the frontal region in Panel A is more relevant for the description of the behaviour of *H. rhodostomus*, than the information from the same region in Panel B for the description of the behaviour of *D. rerio*.

**Figure 4:**
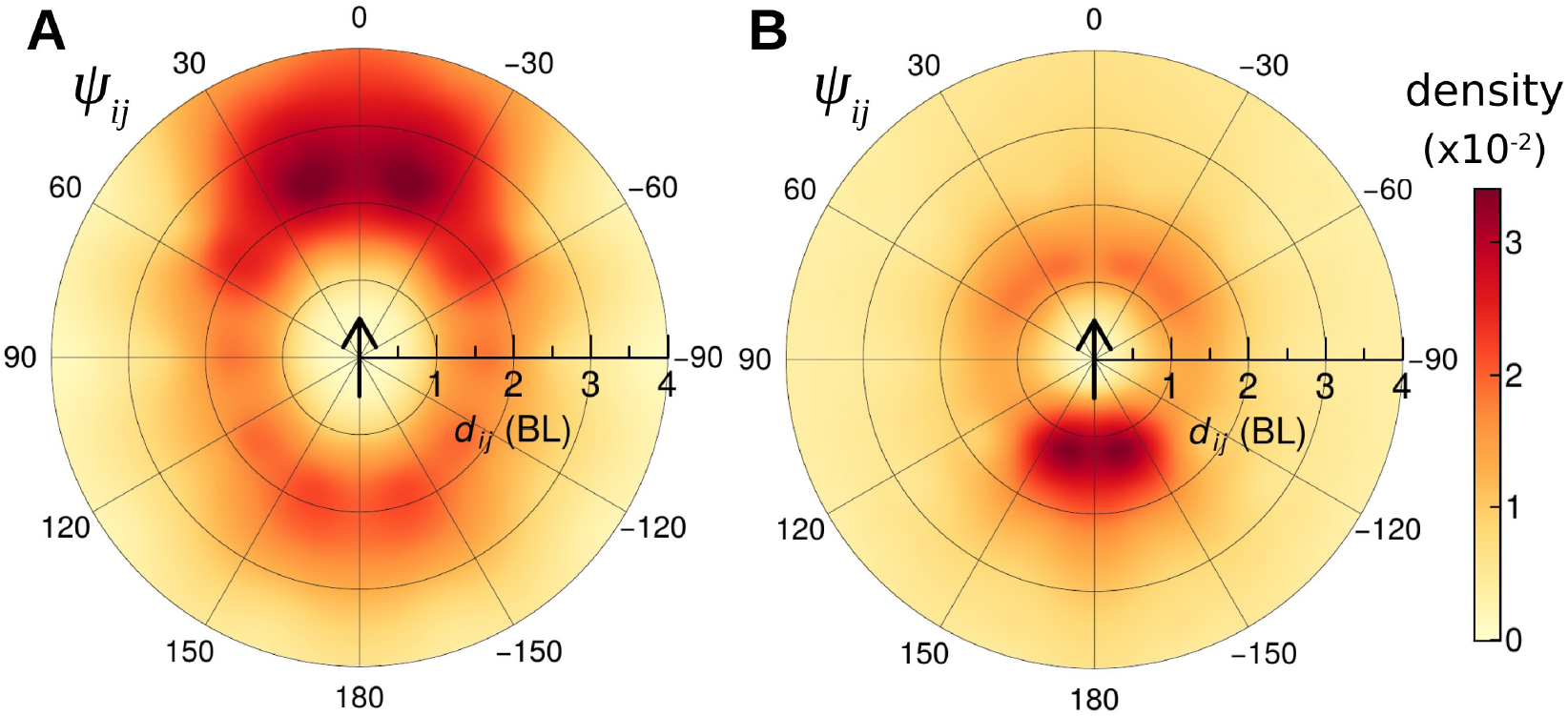
Density maps of the location of a neighbouring fish *j* in the system of reference centred on the focal fish *i*. (A) H. rhodostomus in an arena of radius 0.25 m, and (B) D. rerio in the arena of radius 0.29 m, when the focal fish is at least at 2 BL (6 cm) in *H. rhodostomus* and 1 BL (4.5cm) in *D. rerio* from the wall of the tank. The vertical arrow pointing north located at the centre of the coordinate system indicates the position and orientation of the focal fish *i*.

The simplicity to visualise and interpret data with force maps comes at the cost of important limitations and oversimplifications. Four of the most critical of these limitations are the following:

i. The reduced number of state variables involved in individual behaviour that force maps can handle is limited to 2, at most 3. This limitation leads to the use of projections and average values that can hide crucial features of the phenomenon under study and that, when data are not homogeneously distributed, can produce wrong or inaccurate observations or conclusions.
ii. The difficulty of identifying and disentangling intermediate contributions such as attraction and alignment to behavioural patterns;
iii. The difficulty of finding analytical expressions of the interaction functions in order to implement them in a predictive mathematical model;
iv. The difficulty of distinguishing the effects of variables (*e.g.*, the distance between fish *d*_*ij*_) from the effect of parameters (*e.g.*, the fish body length).

Let us describe these limitations in more detail.

(*i*) When a function depends on more than two variables, force maps are mere projections of the function on a 3D surface, where the value of the function has been averaged with respect to one or more variables. Averaging raises two problems. First, the function can be odd with respect to the variable used to calculate the average, as it is the case of the heading change *δφ* with respect to the angle of perception *ψ*_*ij*_ and the relative heading *φ*_*ij*_ in *H. rhodostomus*, where *δφ*(*d*_*ij*_, *ψ*_*ij*_, *φ*_*ij*_) = −*δφ*(*d*_*ij*_, −*ψ*_*ij*_, *φ*_*ij*_) = −*δφ*(*d*_*ij*_, *ψ*_*ij*_, −*φ*_*ij*_) [12]. Then, for instance, huge variations of *δφ* with respect to *ψ*_*ij*_, but of different sign, can cancel each other and yield a small average value, as if *δφ* was almost independent of *ψ*_*ij*_, so that a crucial feature of the behaviour would be hidden by the averaging process. Second, averaging by simply adding the values and dividing by the number of values implicitly assumes that the probability of occurrence is the same for all the possible states.^1^ This would mean for instance that the probability for a fish *j* of being at the state *d*_*ij*_ = 1 BL, *ψ*_*ij*_ = 90° and *φ*_*ij*_ = 10° with respect to a focal fish *i*, is the same than the probability of being at (*d*_*ij*_, *ψ*_*ij*_, *φ*_*ij*_) = (2, 0, 180), which is clearly not the case, at least in *H. rhodostomus* [12]. Thus, averaging can only provide partial information, can hide crucial effects, and can even produce a completely wrong result.

Adding the map *δφ*(*ψ*_*ij*_, *φ*_*ij*_) to those in Figs. 2AB simply reveals how *δφ* depends on (*ψ*_*ij*_, *φ*_*ij*_), but does not help to understand the relation of these variables with the distance. The visualisation of *δφ* as a function of more than two variables, for instance *d*_*ij*_, *ψ*_*ij*_ and *φ*_*ij*_, would require a large series of maps in which one of the variables is kept constant in each map and varies from one map to another.

As a consequence, the small number of dimensions that can be represented by force maps constitutes a crucial limitation for the description of behaviour, especially if more variables are taken into account, as, *e.g.*, the relative speed v_*ij*_, or the interaction of fish with an obstacle (such as the wall of a tank), which would require two more variables *r*_w,*i*_ and *θ*_w,*i*_, the distance and angle to the wall, respectively. Even for the depicted intermediate values, the precise contributions of each variable on the change of heading are still entangled. For instance, how *φ*_*i*_ varies when *j* points left and is located at the right side of *i*, and how this change varies with the distance *d*_*ij*_? The contribution of each variable cannot be disentangled from those of the other variables.

Moreover, force maps of 3D-functions like *δφ* require huge amounts of data. Using 30 bins per variable with an average of 100 points per bin would require 2.7 × 10^6^ data points, which is very far from being the amount of data collected in most of the experiments on collective motion. We anticipate here that, in order to get the same level of precision, our procedure reduces the required number of data up to 150 times.

(*ii*) Another limitation of force maps is that intermediate contributions are difficult or often impossible to identify. A function *f* can depend on the state variables *x* and *y* through two intermediate functions *a* and *b* such that *f* (*x, y*) = *f* (*a*(*x, y*), *b*(*x, y*)). This is precisely the case of the heading change *δφ*, which depends on the state variables *d*_*ij*_, *ψ*_*ij*_ and *φ*_*ij*_ through the combination of (at least) two intermediate contributions, attraction and alignment, themselves depending on the three state variables [12]. Moreover, attraction and alignment can have opposite contributions, so that their combined effect can be cancelled as if the fish were isolated from each other, while both forces are in action. Force maps are not able to identify which part of the heading change would be due to the attraction or the alignment, that is, the functions *a*(*x, y*) and *b*(*x, y*) cannot be extracted from force maps. Even if representations in 7D were possible, these “maps” would not allow to identify intermediate contributions.

(*iii*) Extracting analytical expressions from a colour map is difficult unless the relation between variables is very simple. However, it is essential to build mathematical or computational models, which will then be used to make predictions in other experimental situations, to draw phase portraits, etc. Simple piecewise linear functions interpolating the data are not suitable because they lack the physical or biological meaning and do not help to build explicit and concise mathematical models. Fig. 2 and 3 show the force maps of *δφ* for two different species of fish. Significant differences appear between maps, *e.g.*, in Panels A, the left-right symmetry of the heading change with respect to the position of the neighbour in *H. rhodostomus*, while in *D. rerio* the symmetry is with respect to the position of the focal fish; in Panels C, the half-planes of homogeneous colour are horizontal in *H. rhodostomus*, but vertical in *D. rerio*. Will such a difference be observed if larger groups are considered? Force maps cannot be extrapolated from one experimental situation to another.

(*iv*) Finally, it is difficult to distinguish the effect of a variable from the effect of a parameter in a force map, *e.g.*, the effect of *d*_*ij*_ from the effect of BL. Are the differences observed in the previous maps of each species due to the physical characteristics of the species (typical body length) or to the experimental conditions (radius of the arena)?

The method we present below does not have these limitations, first, because it can handle a large number of state variables (*i.e.*, of dimensions), and second, because the analytic expressions it provides are precisely the optimal way of disentangling the interaction functions at play and of describing the role of each state variable and each parameter of the system. These analytic expressions can then be exploited in building an explicit and yet concise model, whose agreement with experiment can be tested, and whose predictions can be further investigated experimentally.

## 3 Method to extract and model social interactions from behavioural data

Section 2 shows that force maps are representations of the relation of one quantity (action force, acceleration, heading variation, etc.) as a function of other quantities (relative position, velocity or orientation, angle of perception of another individual, etc.). These quantities only make sense in a framework of the physical world described by a mathematical model which, even if it is not mentioned explicitly in studies [17, 18, 20, 21] (but see also [13, 22, 23]), is usually based on equations of motion built in analogy with Newtonian mechanics.

### 3.1 Outline of the method

Fig. 5 shows the flowchart of the method used to extract and model the social interactions. The method starts with collecting data (step 1) and making a preliminary analysis of the main observables that can be used to describe the phenomenon under study (step 2). In the case considered here, this consists of determining the state variables *d*_*ij*_, *ψ*_*ij*_ and *φ*_*ij*_, and calculating and then analysing a series of measures characterizing the individual behaviour of fish, such as the probability density function (PDF) of the spatial distribution of individuals, their velocity, their distance and orientation to obstacles, their heading variation, and their collective behaviour when swimming in pairs: PDF of the cohesion, which is the distance between individuals, of their relative positions, and of the polarization, which is their relative heading (see Fig. 6). These measures are precisely the state variables identified in Sec. 2. This analysis relies exclusively on the experimental data and is independent of the model.

**Figure 5:**
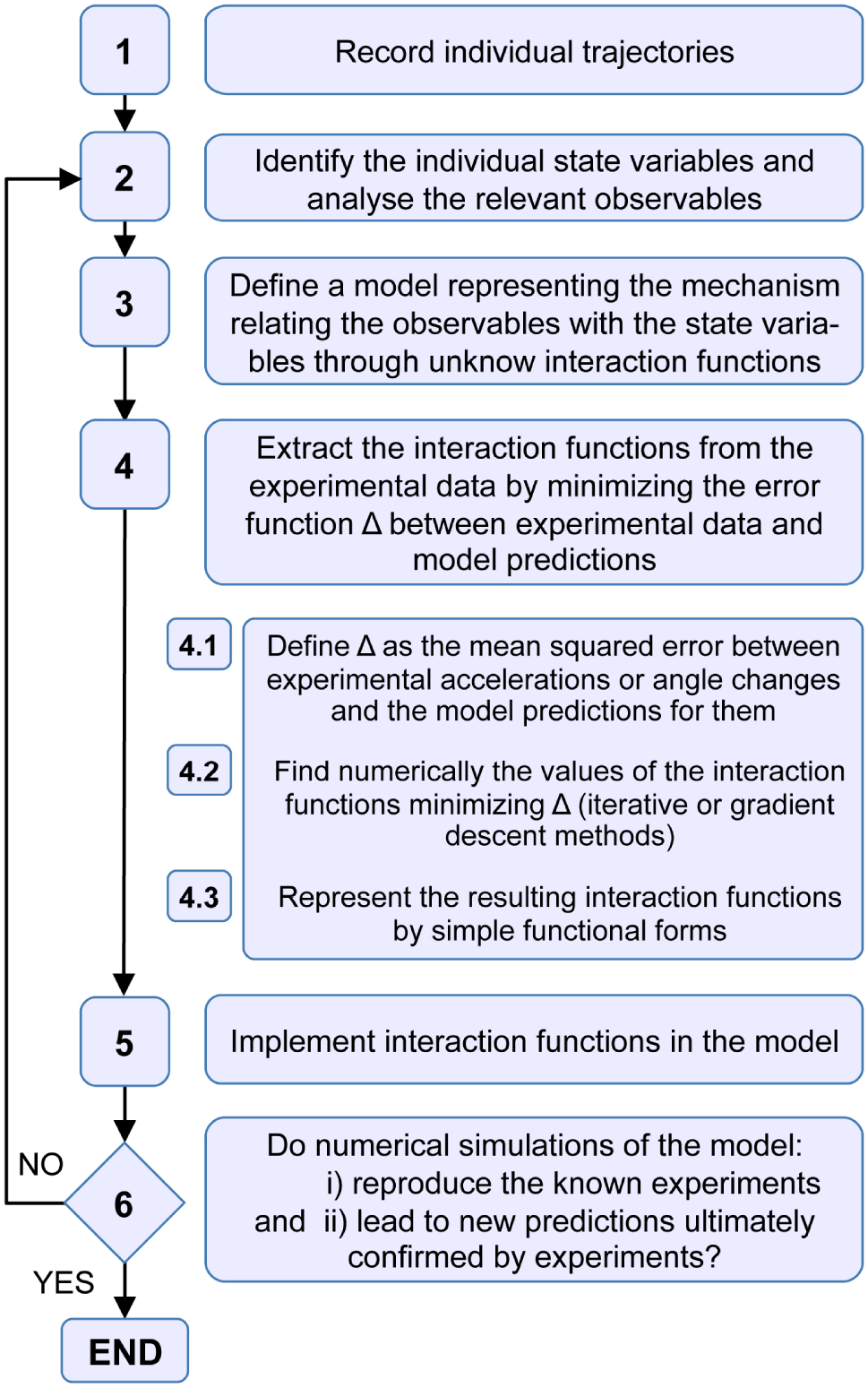
Schematic flowchart of the methodology, with special emphasis on the procedure of extraction of the interaction functions (step 4).

**Figure 6:**
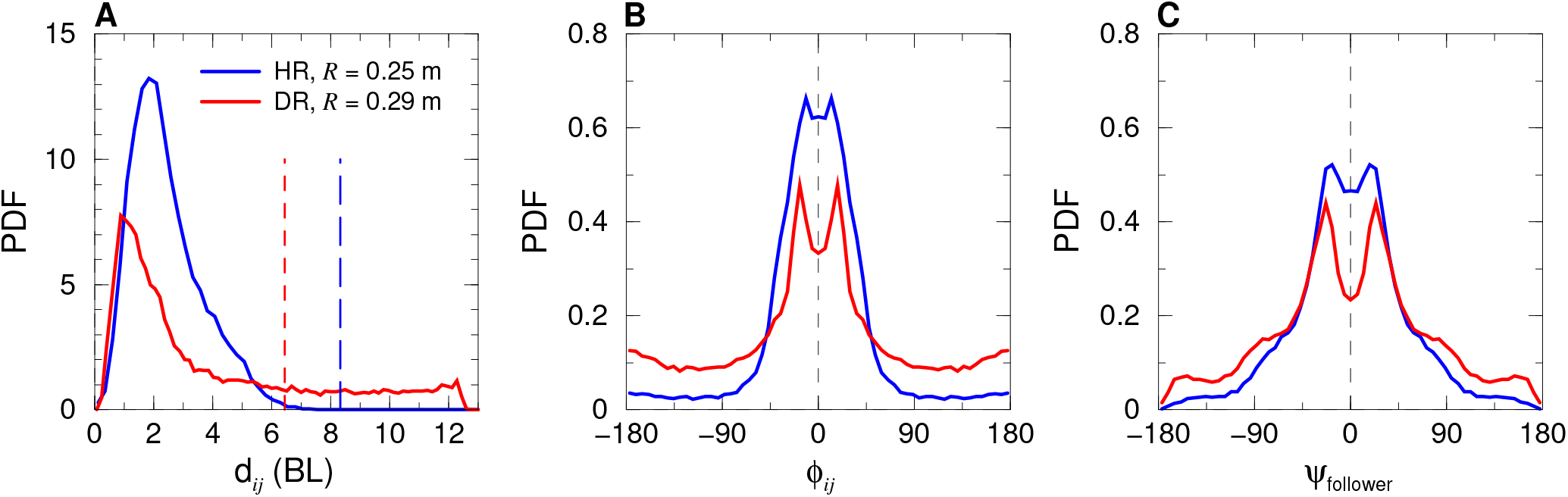
Probability density functions (PDF) of (A) distance between fish *d*_*ij*_, (B) difference of heading *φ*_*ij*_, and (C) angle *ψ*_follower_ with which a fish *i* perceives its neighbour *j* when |*ψ*_*ij*_| < |*ψ*_*ji*_| (the “geometrical follower” is hence the fish which would have to turn the less to face the other fish, the latter being called the “geometrical leader”). Blue lines: *H. rhodostomus* in a circular arena of radius 0.25 m, red lines: *D. rerio* in a circular arena of radius 0.29 m. Vertical dashed lines in panel (A) denote the relative radius *R/*BL of the arena with respect to fish body length of each species, 8.3 in *H. rhodostomus* and 6.4 in *D. rerio* (with 1 BL= 0.03 m in *H. rhodostomus* and 0.045 m in *D. rerio*).

The next step consists in defining a class of models that can provide the more suitable description of the observed phenomenon (step 3 in Fig 5). This is by far the most delicate part of the method. This process has been described in detail in [12] in the case of *H. rhodostomus*; here we simply sketch it, as it is identical for *D. rerio*. We first consider that fish move straight during short time intervals of different duration. These intervals of time are separated by instantaneous “kicks” during which fish adjust their direction and make an abrupt acceleration, after which the speed decays exponentially as the fish glides between two kicks. These assumptions come from the analysis of the trajectories and the variation of the velocity [12]. We also assume that the effects of social interactions occur exclusively when kicks are performed, that is, at the decision times when fish adjust their heading. Then, we provide an explicit equation for the heading variation of the focal fish *δφ* at the moment of a kick, in terms of the quantities that can potentially have an effect on *δφ*.

We now focus on this equation, that accounts for the effects of the social interactions between fish. The selection of the duration and length of the gliding phase and the motion of the fish (with quasi exponential decay of the velocity) between two kicks are given by a simple analytical probability distributions fairly reproducing the experimental ones; this is described in detail in [12]. The equation for the heading variation *δφ* thus reads

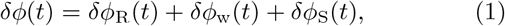

where *δφ*_R_ is a random angle change accounting for the spontaneous decisions of the fish, *δφ*_w_ is due to the repulsion of the wall when the fish is close to a tank wall, and *δφ*_S_ is due to the social interactions with other fish, usually attraction, alignment, or a combination of both, so that

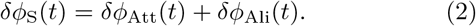

Eqs. (1)–(2) are based on the hypotheses that *δφ*_R_, *δφ*_w_, *δφ*_Att_ and *δφ*_Ali_ are the main contributions to the heading variation of a fish, and that these contributions are combined linearly (additive hypothesis).

Here starts the step 4 of the method, which is the one we wish to emphasize in this article. This step consists in extracting from the experimental data the interaction functions determining the contribution of a neighbouring fish and of the environment to the instantaneous heading variation of a fish.

The interaction of a fish with obstacles such as the wall of an experimental tank is assumed to depend only on its distance *r*_w_ and angle of incidence *θ*_w_ to the wall. The social interactions of a fish depend on the relative state of its neighbours. The state of a fish *j* with respect to a focal fish *i* is given by the distance between them *d*_*ij*_, the relative speed v_*ij*_, the angle with which *j* is perceived by *i*, *ψ*_*ij*_ (≠*ψ*_*ji*_), and the relative heading *φ*_*ij*_. The angle change *δφ*_S_ resulting from social interactions is thus a function of four variables.

Discretising the 6 state variables and averaging as we did in Sec. 2 to build force maps, Eq. (1) becomes

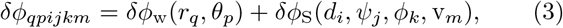

in a grid of *Q × P × I × J × K × M* boxes in 6D. The noise term disappears as it is assumed to have zero mean in each box. If seven-dimensional representations were possible and manageable, we would have behavioural maps of social interactions. But even in that case, the contributions of each variable would still be entangled, and intermediate functions impossible to detect.

For the sake of simplicity, we consider here the case where fish are far enough from the tank wall so that the contribution of *δφ*_w_ to the heading variation can be neglected with respect to the effects of social interactions. The procedure can easily be extended to extract both the social interactions and the interaction with the wall, but to the cost of heavier notations. We refer the reader to Calovi *et al.*’s work [12], where the disentangling of the combined effects of a tank wall and social interactions between fish has been performed.

The key of the procedure of extraction of interaction functions described below consists in assuming that the contribution of each state variable to each kind of interaction can be separated in a product form (which is the case for physical particles [12]):

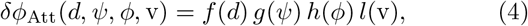

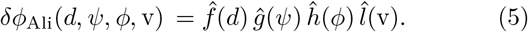

Note that functions with a hat are in general different from the functions without hat, (*i.e.*, 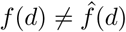, etc.). With this additional hypothesis, the procedure provides analytical expressions of the interaction functions of attraction and alignment and of each contribution to the heading variation. If one of the interactions (attraction or alignment) is not actually at play, the procedure will highlight it. In spite of this separation of variables, possible correlations between magnitudes are in part preserved (see below).

The functions 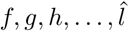 are then discretised and the heading change *δφ*_*ijkm*_ is averaged in 4-dimensional boxes defined by [*d*_*i*_, *d*_*i*+1_] × [*ψ*_*j*_, *ψ*_*j*+1_] × [*φ*_*k*_, *φ*_*k*+1_] × [v_*m*_, v_*m*+1_], so that Eq. (3) becomes

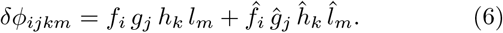

Eq. (6) is a system of *I × J × K × M* equations but involving only only 2(*I* +*J* +*K* +*M*) unknowns. Note that systems with more equations than unknowns are called *overdetermined systems* and rarely have a solution.

The reduction of the system (6) to a solvable one is carried out by minimizing the error 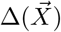 with which this system is satisfied by the set of values for a candidate solution. This is the step 4.1 of the flowchart of our method, shown in Fig. 5. This minimization procedure involving a large number 2(*I* + *J* + *K* + *M*) of variables can be achieved by several methods, and in particular, iterative methods. Importantly, note that the possible correlations existing between the contributions of the different state variables, due to the fact that the system derives from a physical phenomenon, are preserved.

Once the overdetermined system (6) is obtained, the procedure consists in reducing this *unsolvable* system to a solvable one. This corresponds to the step 4.2 of the flowchart in Fig. 5. Box 1 shows how to perform this reduction for a simple 2D-case. The first step consists in writing the error function 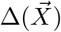, which is a function of *D* = 2(*I* + *J* + *K* + *M*) variables,

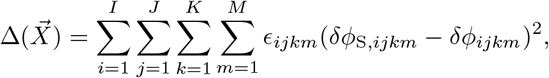

where 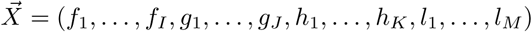 is a vector of ℝ^*D*^ and 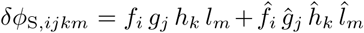 is the discretisation of the social force in each *ijkm*-box. The factor *ϵ*_*ijkm*_ is precisely the number of data in the *ijkm*-box, and serves to preserve the structure of the dataset (by weighting the regions with more data) and the correlations between variables resulting from the dynamics (see also note 1 in Sec. 2).

Following Box 2, we look for the minima of 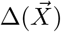 by finding the zeros of the gradient of ∆, 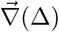. This is a minimization process that can be carried out by different methods (descent methods or other iterative methods). Here we do it by solving the following system of *D* equations for the partial derivatives of ∆:

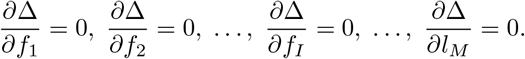

The partial derivative of ∆ with respect to, *e.g.*, *f*_*i*_, is

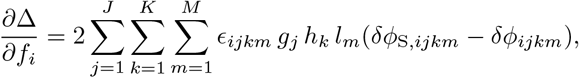

so the equation *∂∆*/*∂f*_*i*_ = 0 of the reduced system is

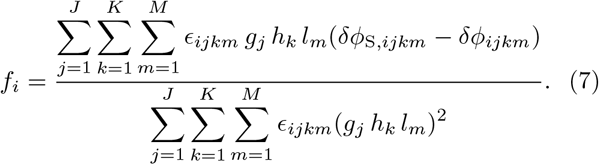

The corresponding equations for the other components have the same structure, where each unknown, as *f*_*i*_ in Eq. (7), is given by an explicit combination of the other unknowns and the known values *δφ*_*ijkm*_ and *ϵ*_*ijkm*_. Note also that *f*_*i′*_, *i*′ ≠ *i*, does not appear in the equation of *f*_*i*_. Thus, the equation of *g*_*j*_ is obtained by replacing “*g*_*j*_” by “*f*_*i*_”, “*j* = 1” by “*i* = 1”, and “*J*” by “*I*” in Eq. (7), and the equation for a function with a hat is obtained by adding a hat to the functions in Eq. (7).

The result is a system of *D* equations and *D* unknowns that can be written in the following form:

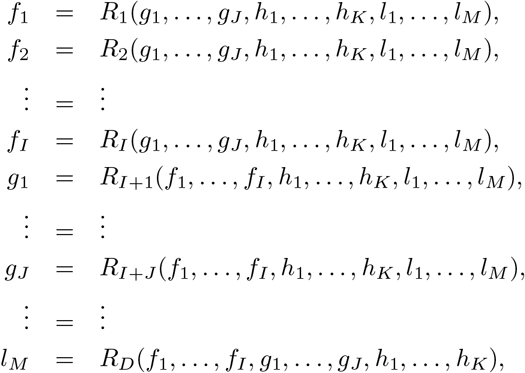

which, in short, is 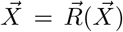, where 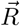 is a function from ℝ^*D*^ to ℝ^*D*^.

Solving such a system amounts to find a fixed point 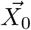 of the function 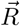. Box 2 shows how to do it by means of an iterative method, and illustrates how this method works in the 1D-case.

In the end, the procedure provides the values of each interaction function at the points representing each box. Depending on the number of boxes, an analytical form of the function can be derived, based on physical principles and specific observations. For example, the function of the angle of perception *g*(*ψ*) must be odd, *i.e.*, *g*(*−ψ*) = − *g*(*ψ*), because the attractive effect of a neighbour on the heading change of a focal fish has the same intensity wherever the neighbour is located at the right or at the left of the focal fish, but has opposite directions in each case: if the neighbour is at the right (resp. left) side, the focal fish would turn right (resp. left) to approach the neighbour. Physical properties of the interactions at play, such as the exponential decay of some interactions, must be taken into account and guide the choice of the final analytical expressions. This part corresponds to the step 4.3 of Fig. 5 and depends on the specific phenomenon under study; it is detailed in Sec. 4 for the case of fish swimming in pairs.

### 3.2 Application to a simple case study

In order to illustrate how the extraction procedure, that is, step 4 of our methodology (Fig. 5) can be used, we have produced artificial data of a simple case in which the interaction functions are known and in which we controlled the level of noise and the distribution of data. This also allows us to illustrate the efficiency of the procedure and the accuracy that can be obtained according to the quality of the data.

Fig. 7 shows the colour map of the following function of two variables *a*(*x, y*): [0, 2] × [−*π, π*] → ℝ,

**Figure 7:**
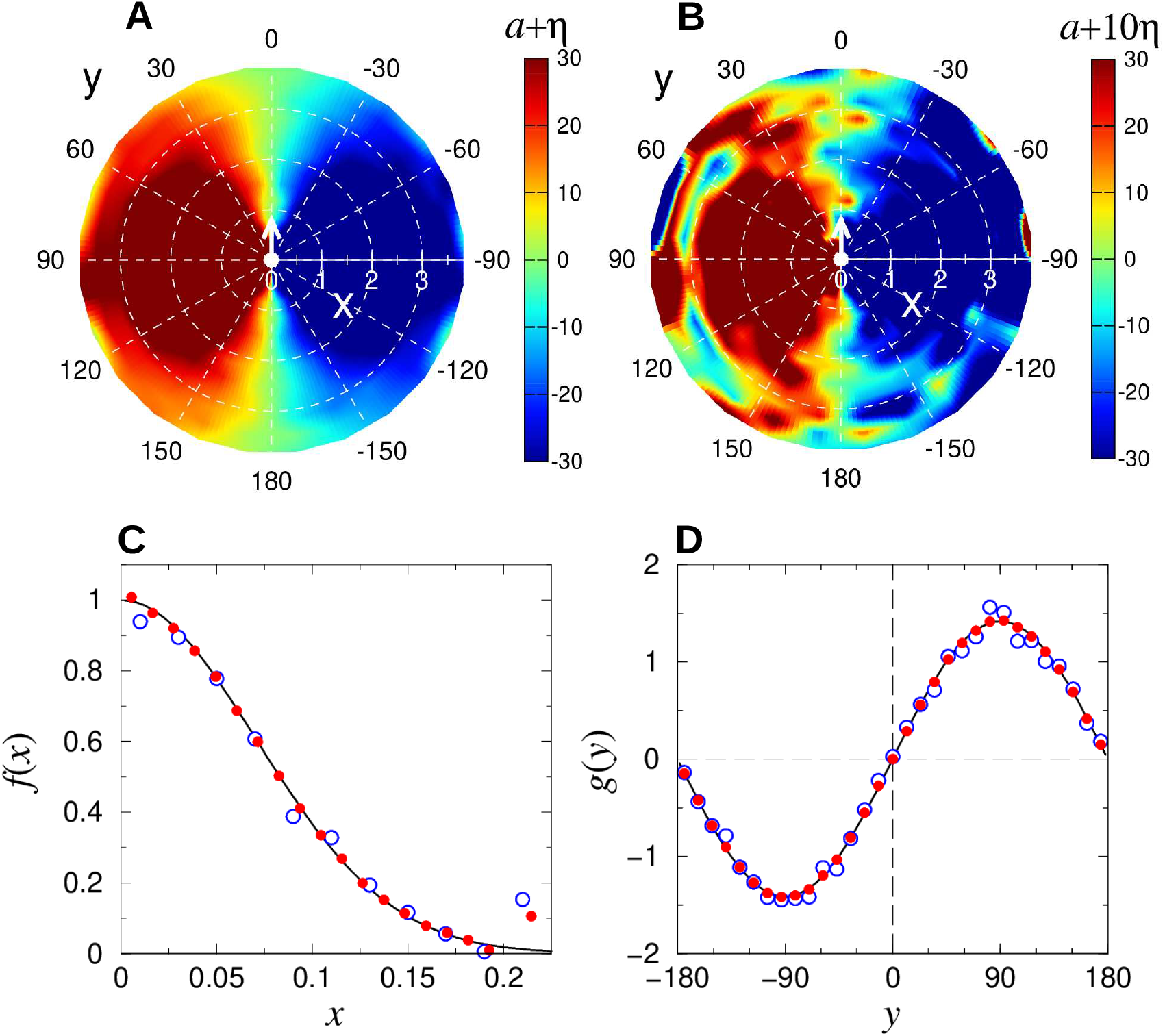
Force maps and reconstruction of the interaction functions in a simple case. (AB) Force maps of the function *a*(*x, y*) + *η*(*x, y*) for two levels of noise, of zero mean and amplitude (A) 0.35, which is the one observed in the experiments of *H. rhodostomus*, and (B) 3.5, a level of noise ten times higher. We used the same distribution of points *ϵ*_*ij*_ than the one observed in the experiments of *H. rhodostomus*. (CD) Reconstruction of the interaction functions (C) *f* (*x*) and (D) *g*(*y*) extracted with our procedure (red dots and blue circles), compared to the analytical expressions exp[−(*x/x*_0_)^2^] and 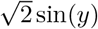 respectively (black lines). Red dots correspond to the case of a noise of amplitude 0.35, where we used 20 nodes in *x* and 31 in *y*, and blue circles to the case of a much higher level of noise (3.5) and with less nodes in *x*, 11 instead of 20. Note that the function *g*(*y*) is normalized.

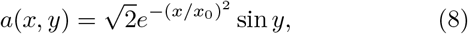

with *x*_0_ = 0.1. The map is built as follows. We first discretise the 2D-space (*x, y*) in *N* × *M* rectangular cells 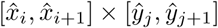, where

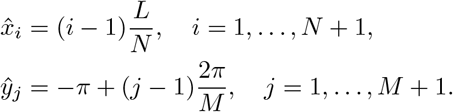

Then, we evaluate the function *a*(*x, y*) on 100000 points randomly selected, and place each point on the corresponding cell. After that, we count the number of points in each cell, *ϵ*_*ij*_, and we assign to *a*_*ij*_ the average of the values of the function for all the points that are in the cell *ij*. To simulate the effect of the noise, which is always present in real data sets, for each point found on cell *ij*, we add a small noise of zero mean and standard deviation = 0.35 to the value of *a*_*ij*_. The resulting value is then considered as the *measured* value of the function *a*(*x, y*) in the corresponding cell, denoted by (*x*_*i*_, *y*_*j*_), and usually defined as the middle point of the cell: 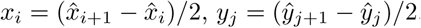.

We now apply the procedure described above, which will lead us to analytical expressions of the interaction functions of *x* and *y* that give rise to this colour map.

In step 3 of our method (Fig. 5), we make the assumption that two decoupled functions *f* (*x*), *g*(*y*) exist such that *a*(*x, y*) = *f* (*x*) *g*(*y*). Evaluating this equation in each cell *ij*, this means that *a*_*ij*_ = *f*_*i*_ *g*_*j*_ for *i* = 1, *…N* and *j* = 1, …, *M*. The value of *a*_*ij*_ is known for all *ij* because it is the mean value of the data found in the cell *ij*, used to build the colour map of Fig. 7. In turn, the values of *f*_*i*_ and *g*_*j*_ are unknown for all *i* and all *j*. These *N × M* equations and *N* + *M* unknowns constitute precisely the overdetermined system (6). Following step 4.1 in Fig. 5, the error function ∆ is written, resulting to be identical to the one shown in Eq. (13) of Box 1. The overdetermined system is thus reduced to the solvable one shown in (17) which, following Box 2, is solved with an iterative method (step 4.2). Figs. 7BC show the points 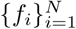 and 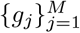 that solve the reduced system (17), in good agreement with the original data despite the addition of noise (including in the case of a much larger noise than the one inferred in the actual data of our fish experiments).

Note that if *f* and *g* are a solution of (17), then the functions (1/*α*)*f* and *αg*, where *α* is a real number, are also a solution of (17), since the product of the two functions remains invariant. Hence, we need to impose an additional condition in order to account for this under-determination and to generally allow for a proper comparison of reconstructed interaction functions. Following [12], we chose to normalize all angular functions such that their squared average is unity. In particular, *g* is normalized such that 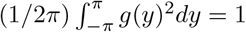, a normalization applied in Figs. 7BC.

The last step consists in finding simple analytical expressions that interpolate the discrete values of the reconstructed interaction functions, in order to implement them in an explicit mathematical/computational model. There is an infinite number of combinations, so that one must be guided by key physical features of the phenomenon under study that are well established, properties such as symmetries, analogies with other physical systems. For example, for angular functions, the parity is often easily identifiable from the data or can be asserted from general principles (mirror symmetry, left/right symmetry...), so that few Fourier modes can be sufficient to interpolate angular functions resulting from the reconstruction procedure.

## 4 Extraction and comparison of social interactions in different species of fish

The model proposed in Sec. 3 is based on the assumption that social interactions are combined in an additive form (see step 3, Fig. 5). In Eq. (2), two functions *δφ*_Att_ and *δφ*_Ali_ were introduced to account for the attraction and alignment interactions, for which no *a priori* assumptions were made except that their dependence on the state variables *d*_*ij*_, *ψ*_*ij*_ and *φ*_*ij*_ is decoupled; see Eqs. (4)–(5). Steps 4.1 to 4.3 in Fig. 5 provide us with functional forms, but do not determine which functional form corresponds to which kind of interaction.

To do that, observations made in Sec. 2 (step 2 of the flowchart in Fig. 5) are used to say that *δφ* must change sign when *ψ*_*ij*_ and *φ*_*ij*_ both change sign, and that, consequently, the same must happen for *δφ*_Att_ and *δφ*_Ali_. This way, the parity of the angular components of the interaction functions is univocally determined. Thus, to have an interaction of attraction, the fish must turn left if its neighbour is on its left, and turn right if it is on its right; that is, *δφ >* 0 if *ψ*_*ij*_ *>* 0 and *δφ <* 0 if *ψ*_*ij*_ *<* 0. Assuming perfect left/right symetry, this exactly means that *δφ*_Att_ must be an odd function of *ψ*_*ij*_, and thus an even function of *φ*_*ij*_, provided fish do not have side preferences to turn (that is, fish do not *prefer* turning left to turning right). Similarly, to have an interaction of alignment, the fish must turn left when the relative heading of its neighbour is turned to the left, and turn right if it is turned to the right; that is, *δφ*_Ali_ must be an odd function of *φ*_*ij*_, and thus an even function of *ψ*_*ij*_.

This allows us to rewrite the interaction functions from Eqs. (4)–(5) as follows:

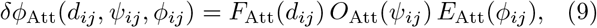

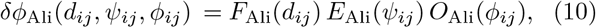

where function names “*O*” and “*E*” stand for “odd” and “even” respectively.

Fig. 8 shows the social interaction functions reconstructed with the procedure described in the previous section in the case of two *H. rhodostomus* (Panels ABC) and two *D. rerio* (Panels DEF) swimming in circular arenas. Attraction and alignment have been detected in both species and, although some important differences can be observed, each kind of interaction has essentially a similar shape and intensity in both species. Hence, we used the analytical expressions introduced in [12] for *H. rhodostomus* (solid lines in Fig. 8) to fit the discrete values of *D. rerio* extracted from the experimental data at Step 4 of our method (Fig. 5), that is, with the procedure described in Section 3 (points in Fig. 8).

**Figure 8:**
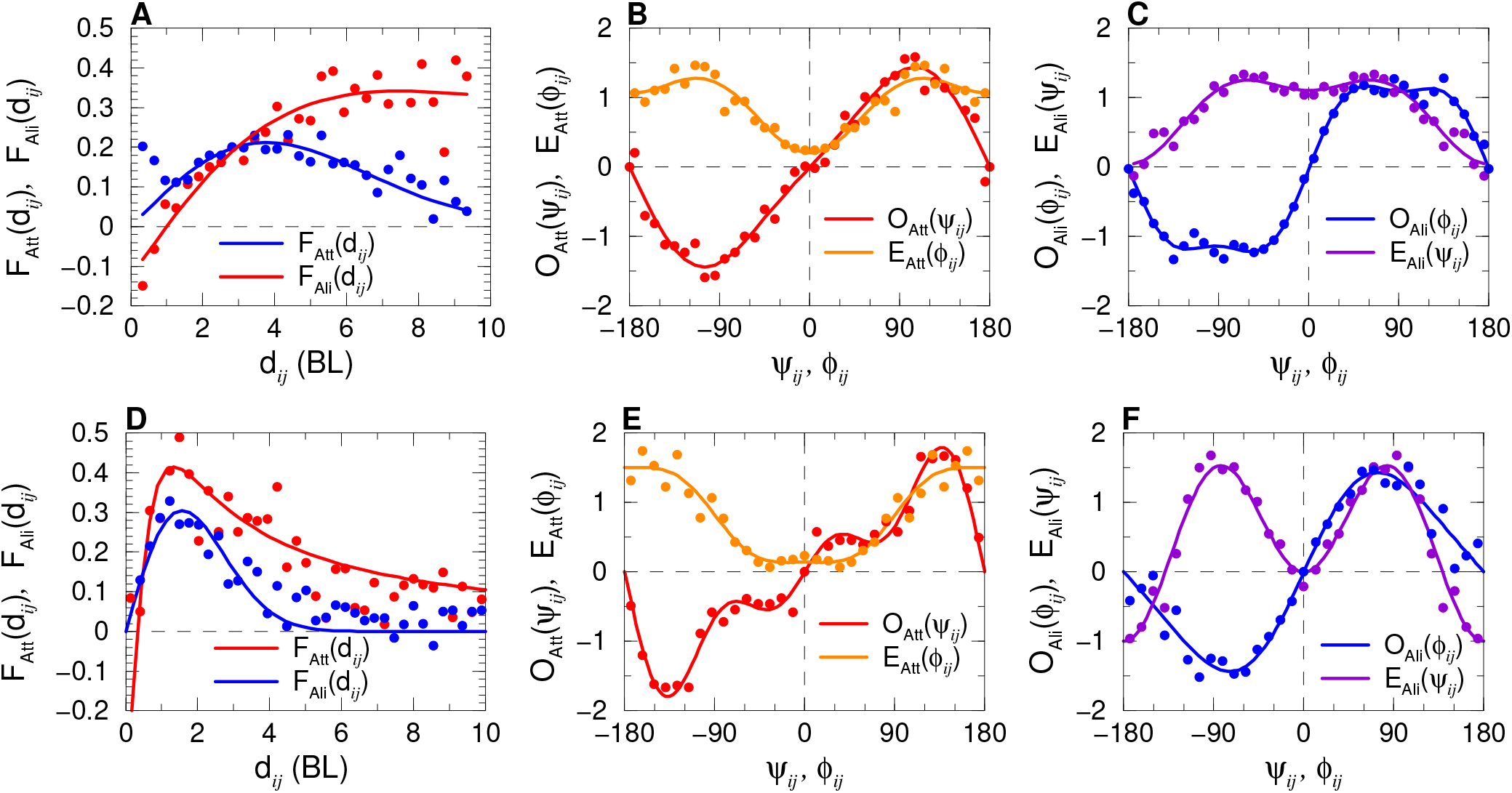
Analytical expressions of the social interaction functions of *H. rhodostomus* (ABC) and *D. rerio* (DEF) swimming in arenas of radius 0.25 and 0.29 m respectively (solid lines), interpolating the discrete values extracted from the experimental data with the procedure described in Sec. 3 (dots). (AD) Intensity of the attraction (red) and alignment (blue), modulated by the angular functions of (BE) attraction and (CF) alignment.

These expressions are, for the intensity of the social interaction of attraction and alignment, as follows:

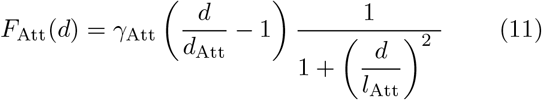

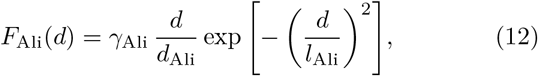

where the values of the parameters depend on the species and on the size of the arena. Note that the expression of the intensity of the alignment has been simplified with respect to the one obtained in [12] and now has one parameter less. Here, *γ*_Att_ and *γ*_Ali_ are the (dimensionless) intensities of the attraction and alignment interactions, *d*_Att_ is the distance below which attraction changes sign and becomes repulsion, *l*_Att_ and *l*_Ali_ are the ranges of each interaction (the higher the value, the longer the range). *d*_Ali_ = 1 BL is used to make *γ*_Ali_ dimensionless and allow direct comparison with *γ*_Att_ and with the intensify of the interaction with obstacles *δφ*_w_ and the free decision term *δφ*_R_ in Eq. (1). Table 1 shows the parameter values corresponding to each species. Note that the values for *H. rhodostomus* have been adapted from those in [12], according to the slightly different expression used here.

**Table 1.** Parameter values for *H. rhodostomus* and *D. rerio* in circular arenas of radius 0.25 m and 0.29 m respectively.

|  | *H. rhodostomus* | *D. rerio* |
| --- | --- | --- |
| $R$ (m) | 0.25 | 0.29 |
| $R/\text{BL}$ | 8.33 | 6.44 |
| $\gamma_{\text{Att}}$ | 0.124 | 0.42 |
| $d_{\text{Att}}$ (m) | 0.03 | 0.015 |
| $l_{\text{Att}}$ (m) | 0.193 | 0.042 |
| $\gamma_{\text{Ali}}$ | 0.092 | 0.32 |
| $d_{\text{Ali}}$ (m) | 0.03 | 0.045 |
| $l_{\text{Ali}}$ (m) | 0.16 | 0.1 |

Regarding the normalized angular functions for *D. rerio*, we found the following expansions (with at most 3 Fourier modes):

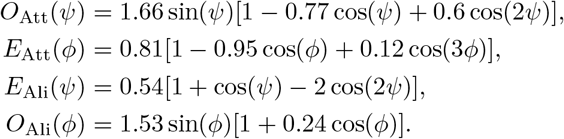

Following the step 5 of the flowchart of our methodology (Fig. 5), these analytical expressions should be implemented in the model introduced in Section 3. Then, numerical simulations of the model should be performed and compared with the known experiments, and predictions should be made, that must be verified *a posteriori*. This is the step 6 of our methodology (Fig. 5); it was done for *H. rhodostomus* in [12], and will be done elsewhere for *D. rerio* and other species.

Comparing the interaction functions found for both species, we observe that the functions are similar in shape, especially the angular functions (see *e.g.* the angular functions of attraction in Fig. 8B and Fig. 6C, for which the same function of the angle of perception *ψ*_*ij*_ could have been used). Both attraction and alignment have intensities of the same order of magnitude in both species: the maximum of the attraction is around 0.4 in both species, and the maximum of the alignment is around 0.2–0.3; see Fig. 8A and Fig. 6B.

The most important difference is that the range of the interactions is between three and four times larger in *H. rhodostomus* than in *D. rerio*: in *H. rhodostomus*, the maximum intensity of the attraction is reached at around 4–6 BL and the one of the alignment at around 2–3 BL, while in *D. rerio* these maxima are at 1 and 1.5 BL for attraction and alignment respectively. Moreover, the intensities decay more rapidly in *D. rerio* than in *H. rhodostomus*, especially with respect to the respective fish body length (alignment intensity is zero beyond 5 BL ≈ 22.5 cm in *D. rerio*, but is still perceivable at 10 BL ≈ 30 cm in *H. rhodostomus*). In *H. rhodostomus*, repulsion acts when the other fish is at less than 1 BL, but only when it is at less than 0.5 BL in *D. rerio*. Attraction almost always dominates alignment in *D. rerio* (the intersection of the red and blue lines in Fig. 8A is at around 0.5 BL), while, in *H. rhodostomus*, alignment was found to dominate attraction at short distances (under 2.5 BL).

In *H. rhodostomus*, a fish *i* is subject to a stronger attraction when the other fish *j* is at its right or left side (highest values of *O*_Att_ are reached when *ψ*_*ij*_ ≈ ±90°, see Fig. 6C) and moves more or less perpendicular to it (*E*_Att_ is higher at *φ*_*ij*_ ≈ 90–100°). Alignment is stronger when the other fish is in front (*E*_Ali_ is higher at |*ψ*_*ij*_| *<* 80°). In *D. rerio*, attraction is stronger when the other fish is clearly behind the focal fish (*O*_Att_ is higher at *ψ*_*ij*_ ≈ 135°) and moves in the opposite direction (*E*_Att_ is higher when |*φ*_*ij*_| *>* 100°; see Fig. 8B).

In both species, the strength of the interaction of alignment vanishes when fish are already aligned (*O*_Ali_ ≈ 0 when |*φ*_*ij*_| *<* 30°. See Fig. 8C and Fig. 6D. The strength of alignment is active in *D. rerio* essentially when both fish are perpendicular to each other (high intensity of |*O*_Ali_| ≈ 1.5 is reached when *φ*_*ij*_ ≈ ±85°) and the focal fish has its neighbour at one of its sides (*E*_Ali_ is peaked at *ψ*_*ij*_ ≈ ±85°), while in *H. rhodostomus*, the ranges of interaction in both the angle of relative heading and the angle of location of the neighbour are much wider: alignment is active when the neighbour is ahead of the focal fish (*ψ*_*ij*_ ∈ (−100°, 100°)) and fish are simply slightly aligned (|*φ*_*ij*_| ∈ (45°, 135°)).

In summary, when swimming in pairs, *H. rhodostomus* interact in a much wider range of situations than *D. rerio*. This is true with respect to the three state variables of a focal fish: 1) in *H. rhodostomus*, the distance *d*_*ij*_ at which both attraction and alignment forces are active is larger than in *D. rerio*; 2) the location of the neighbour where the forces are active is much wider in *H. rhodostomus* than in *D. rerio*, and 3) the same happens for the range of relative headings. The maximum intensity of the interaction is similar in both species, although high values of the intensity are reached in a much wider range of situations in *H. rhodostomus* than in *D. rerio*.

## 5 Discussion and conclusions

Behavioural biology has recently become a “big-data science” mainly supported by the advances in imaging and tracking techniques. These new tools have revolutionized the observation and quantification of individual and collective animal behaviour, improving to unprecedented levels the variety and precision of available data [9, 24, 25, 26]. As access to large volumes of data is gradually stepping animal behaviour research into a new era, there is also a growing need for understanding interactions between individuals and the collective properties that emerge from these interactions. Animal societies are complex systems whose properties are not only qualitatively different from those of their individual members, but whose behaviours are impossible to predict from a prior knowledge of individuals [3, 4]. However, understanding how the interactions between individuals in swarms of insects, schools of fish, flocks of birds, herds of ungulates or human crowds give rise to the “collective level” properties requires the development of mathematical models. These models allow to investigate of how complex processes are connected, to systematically analyse the impact of perturbations on collective behaviour (*e.g.*, when a predator is detected in the neighbourhood), to develop hypotheses to guide the design of new experimental tests, and ultimately, to assess how each biological variable contributes to the emergent group level properties.

We have presented a general methodology which leads to the measurement of the social interactions (defined in the framework of a model structure) from a set of one-individual and two-individual trajectories. This procedure, illustrated here on two different fish species, can be similarly applied on any set of trajectories of other organisms, including humans [27, 28]. Once the experimental trajectories have been obtained, the extraction of interaction functions only makes sense after they have been defined in the framework of a general model for the equation of motion of an individual interacting with its environment (obstacles and another individual). The model should involve the relevant variables regarding the interaction with obstacles (distance to the nearest wall, angle of the velocity with respect to the normal to the wall...) and another individual (distance between individuals, relative velocities, viewing angle...). In addition, each interaction components (repulsion, attraction, alignment...) is reasonably assumed to contribute additively: for instance, the influence of the wall and another individual on the focal individual is the independent sum of the two corresponding interactions. More importantly, the central simplification in our approach consists in assuming that each interaction contribution can be adequately described by a product of (yet unknown) single-variable interaction functions of the relevant variables (or of a combination of these variables). The structure of the model is obviously also constrained by the considered species, their motion mode, and their anticipated interactions. For instance, for fish displaying a burst-and-coast swimming mode, the dynamical model is intrinsically discrete in time and returns the angle change of an individual after each kick. For humans or some other fish species with a smooth swimming mode, a continuous-time model is necessary.

The unknown interaction functions defined in the structure of the model are not constrained, and the aim of the extraction procedure (step 4 in Fig. 5) is to measure them, without any a priori assumption about their form or intensity. In order to achieve that, each unknown single-variable interaction function is tabulated on a one-dimensional grid, and their values at each grid point are the fitting parameters. These parameters are then determined by minimizing the mean quadratic error between the prediction of the model and the experimental angle changes after a kick (discrete dynamics) or the experimental acceleration (continuous time dynamics). This minimization process (Box 1) is achieved by solving, for instance with an iterative method (Box 2), the equations expressing the vanishing of the partial derivatives of the error with respect to each fitting parameters, as we have done here, or by gradient descent methods. Once the interaction functions have been obtained on their respective grid, they are fitted and represented by simple analytical forms capturing at best their general shape (Fig. 8).

The original general equation of motion is now complemented by these explicit interaction functions, leading to a concise and explicit model, which can be straightforwardly implemented numerically and can even be studied mathematically. The ability of the model to reproduce the experimental results and even to predict the behaviour of individuals in other situations not yet investigated experimentally can then be assessed (the original model can be extended and the full extraction procedure repeated if an important feature appears to be missing). In particular, quantities like the probability distributions of the distance to the wall, of the distance between two individuals, of the angle between the two velocities or the velocity and the normal to the wall, as well as other observables, permit to assess the predictive power of the model. Note that the interaction functions appearing in the model are determined by finding the best equation of motion describing the instantaneous decisions of the individuals. It is by no means trivial that this is enough for the resulting dynamical model to be able to reproduce observable quantities measured after averaging over many trajectories, or to predict the behaviour of individuals in different experimental conditions. A model able to achieve this certainly provides a convincing indication that its original and general structure and its extracted interaction functions properly represent and describe the behaviour and motion of the considered species.

The main limitations of our methodology lie in the reasonable assumption of additive contributions for the different interactions, and more critically, in assuming that each of these interactions is the product of single-variable interaction functions. This is the cost to pay for only involving a limited number of fitting parameters, yet capturing a large part of the complex structure of these interactions, and also for ultimately obtaining a concise, explicit, and exploitable model.

To address these issues, we are planning in the near future to test our models on a robotic platform [29]. Such a platform could also allow us to study bidirectional interactions between robots capable of reproducing the trajectories generated by our models and real fish, to obtain a more representative validation of our models of interactions [30].

Yet, the advantages and benefits of our approach are numerous. First, the number of fitting parameters (typically 30 for each of the typically 5-8 interaction functions), although apparently large (a total of typically 150-300 parameters), is in general much smaller than the number of data points in the available experimental trajectories. In comparison, a complete force map in typically 5 or more dimensions (one dimension per relevant variable) on a mesh involving 30^5^ boxes, with enough data points in each of them, would require millions if not billions of experimental data points, which is in general impossible to achieve. Two-dimensional force maps obtained after projection (*i.e.*, averaging on the other 5 − 2 = 3 variables), although less noisy than the original force map, cannot be exploited to build a model, since it can be shown that they are strongly affected by the existing correlations between these variables along actual trajectories. For instance, if a fish is very close to a wall, there is a high probability that the fish is in fact parallel to the wall, so that its distance from the wall is strongly correlated to its heading angle. On the other hand, it can be shown [12] that our methodology to extract interaction functions is not affected at all by the likely correlations present in the system, and actually exploit them.

Our method is also robust with respect to the presence of noise in the data (intrinsic behavioural noise or unwanted experimental noise), and can actually characterize the spontaneous fluctuations of the speed and heading angle of the individuals [12, 28]. Moreover, the extraction of interaction functions requires very limited computing power, being obtained in a few seconds on a standard workstation. Ultimately, the analysis of the resulting interaction function allows to make general qualitative conclusions about the interaction at play for the considered species. Force maps can also help in this analysis, but our approach allows for an even finer analysis thanks to the disentangling of interactions by means of separate and explicit interaction functions, instead of mere projected colour maps affected in an unknown manner by the inherent correlations present in the system and the mixing of the different interaction contributions.

More importantly, our methodology ultimately leads to a concise and explicit model which can be exploited to understand and explain diverse experimental features and various forms of collective behaviour, and which presents a predictive power, while force maps cannot be directly exploited to build such explicit models. Note that the structure of the model should be robust for different species having comparable motion mode. For instance, this structure is the same for *H. rhodostomus* and *D. rerio* (or for any species with a burst-and-coast or run-and-tumble motion mode), and the behavioural differences between the species is solely and fully encoded in their different measured interaction functions.

In the specific case of *H. rhodostomus* and *D. rerio*, we have also found that the interaction functions characterizing their interaction with the wall are very similar (see [12] for *H. rhodostomus*), which explains their common tendency to swim close to the wall, especially for isolated individuals. However, even if the interaction functions describing the repulsion/attraction and alignment interactions between two fish have a similar general structure and shape for both species (Fig. 8), the range of the attraction and alignment interactions is much shorter for *D. rerio*. In addition, the intensity of both interactions for *D. rerio* is strongly reduced when the focal fish is behind and hence follows the other fish (|*ψ*_*ij*_| *<* 30°). Both features contribute to a much weaker coordination of the motion in groups of two fish in *D. rerio*, compared to *H. rhodostomus*, which can already be qualitatively noticed by observing recorded trajectories. This weaker coordination is quantitatively illustrated in the measured probability distribution of the distance between two fish (Fig. 4A), which is wider for *D. rerio* and extends up to much larger distances, and of the heading angle difference between the two fish (Fig. 4B), less peaked near *φ*_*ij*_ = 0 for *D. rerio*. Finally, the weaker coordination for *D. rerio* translates in a similar behaviour for the (geometrical) leader and follower, whereas the leader and follower have very distinct behaviour for *H. rhodostomus* (both quantitatively reproduced by the model [12]).

### Materials and methods

#### Ethics

Experiments with *H. rhodostomus* have been approved by the Ethics Committee for Animal Experimentation of the Toulouse Research Federation in Biology No. 1 and comply with the European legislation for animal welfare. Experiments with *D. rerio* were conducted under authorization approved by the state ethical board of the Department of Consumer and Veterinary Affairs of the Canton de Vaud (SCAV) of Switzerland (authorization No. 2778). During the experiments, no mortality occurred.

#### Study species

*H. rhodostomus* were purchased from Amazonie Lab`ege (http://www.amazonie.com) in Toulouse, France. Fish were kept in 150L aquariums on a 12:12 hour, dark:light photoperiod, at 26.8°C (±1.6°C) and were fed *ad libitum* with fish flakes. The average body length of the fish used in the experiments was 31 mm. Wild-type *D. rerio* with short fins (AB strain) were acquired in a number of 60 from a pet shop, and stored in a 60-litre aquarium. The average body length of the fish used in the experiments was approximately 4.5 cm in length. The water in the housing aquarium was kept at a temperature of 26°C. The fish were fed once per day with commercial food between 16:00 and 18:00. Additionally, we opted to use enrichment for the aquarium in the form of plastic plants, Cladophora, gravel, rocks, and aquatic snails.

#### Experimental procedures and data collection

The experimental tank (120 × 120 cm) used to investigate swimming behaviour in *H. rhodostomus* was made of glass and was set on top of a box to isolate fish from vibrations. The setup, placed in a chamber made by four opaque white curtains, was surrounded by four LED light panels giving an isotropic lighting. A circular tank of radius *R* = 25 cm was set inside the experimental tank filled with 7 cm of water of controlled quality (50% of water purified by reverse osmosis and 50% of water treated by activated carbon) heated at 26.1°C (±0.3°C). Reflections of light due to the bottom of the experimental tank are avoided thanks to a white PVC layer. Each trial started by setting two fish randomly sampled from their breeding tank into a circular tank. Fish were let for 10 minutes to habituate before the start of the trial. A trial consisted in one or three hours of fish freely swimming (*i.e.*, without any external perturbation) in a circular tank. A total of 16 trials were performed. Fish trajectories were recorded by a Sony HandyCam HD camera filming from above the set-up at 50 Hz (50 frames per second) in HDTV resolution (1920×1080p).

In the case of *D. rerio*, the experimental setup had dimensions of 100×100×25 cm^3^, inside which a circular tank of radius *R* = 29 cm was placed. The sides and bottom part of the tank were covered with a Teflon plate to avoid reflections. Furthermore, the setup was confined behind white sheets to isolate the fish from external stimuli in the room, while also maintaining a consistent lighting environment inside the setup bounds. A uniform luminosity for the room was provided by four 110 watt fluorescent lamps placed at each of the four sides of the tank. Prior to placing fish in the experimental setup, we ensured that the height of the water was 6 cm. 10 videos of 70 mn duration each were recorded in the circular tank. Subsequently, a group of fish was randomly selected and caught from the rearing tanks to participate in the experiment. A pair of fish was then chosen and placed in the setup. We allowed the fish to habituate for 5 mn before starting the 70 mn long recording. After a single experiment was completed, the fish were returned to the original rearing tank without being re-inserted in the selected group (*i.e.*, no individual was used twice in the same day). The positions of fish on each frame were tracked with idTracker 2.1 [8]. Time series of positions were converted from pixels to meters and the origin of the coordinate system was set to the centre of the ring-shaped tank. Tracking errors (approx. 20% of the data) were corrected and instances where at least one fish moves less than 0.5 body length per second during 4 seconds were removed. More than 8 hours remained during which one fish can kick.

#### Segmentation of trajectories

Instants of kicks were identified as local minima of the velocity preceding local maxima (which are more easy to identify), along time intervals [*t − t*_*w*_, *t* + *t*_*w*_], where *t*_*w*_ is a time window of 0.32 seconds. Trajectories are thus considered as a sequence of kicks. Fish almost never kick at the same time (asynchronous kicks); the position of the other fish is calculated at each instant of kick of the focal fish by simple linear interpolation.

## Acknowledgements

This work was supported by the Germaine de Staël project No. 2019-17 and the Swiss National Science Foundation project “Self-Adaptive Mixed Societies of Animals and Robots”, Grant No. 175731. R. E. was supported by Marie Curie Core/Program Grant Funding, Grant No. 655235–SmartMass. V.L. was supported by doctoral fellowships from the scientific council of the Université Paul Sabatier. We thank Gérard Latil for technical assistance.

### Box 1. Reduction of an overdetermined system of equations

Usually, overdetermined systems do not have a solution, unless the equations have some relation between them (mathematically, if the surplus equations are linear combinations of the others). When equations derive from data extracted from a real phenomenon, a relation is expected to exist between them, based on the underlying laws governing the phenomenon under study. However, finding the solution directly is still difficult, and often plagued by experimental noise, so that the efforts are spent in finding a set of values that minimizes some norm of the error, usually the *l*_2_-norm (as, *e.g.*, in the least squares method). For example, the squared *l*_2_-norm of a vector of two components is 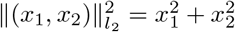.

In a system of the form *x*_*i*_*y*_*j*_ = *a*_*ij*_, *i* = 1, …, *N*, *j* = 1, …, *M*, which is the form of the system (6), the error in each equation is given by |*x*_*i*_*y*_*j*_ − *a*_*ij*_|, and its *l*_2_-norm is the function ∆:

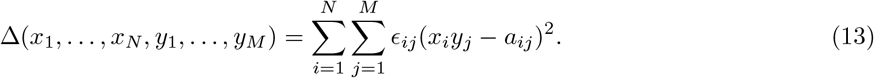

Here *ϵ*_*ij*_ is the number of data that fall in the cell *ij*. When *ϵ*_*ij*_ is the same for all *ij*, the function ∆ is exactly proportional to the *l*_2_-norm. However, for systems that derive from physical and biological phenomena, it is fundamental to modulate the contribution of each cell, not only to give a higher weight to the more frequent values, but also to preserve actual correlations that can exist between the variables.

The goal is thus to find a set of values *x*_1_, …, *x*_*N*_, *y*_1_, …, *y*_*M*_ that minimizes ∆. ∆ = 0 implies that *x*_*i*_*y*_*j*_ = *a*_*ij*_, for all *i* and *j*, and that the system is solved exactly. Finding the minima of a function consists in finding the zeros of its derivative, which, in several dimensions, is the *gradient vector* 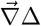, given by the partial derivatives of ∆ with respect to its components; see the Box *Finding minima* below.

##### Finding minima

A minimum of a function *f* is a point *x*_*m*_ where the derivative of the function *f*′ is zero and the values of *f* around the point *x*_*m*_ are higher than *f* (*x*_*m*_). Note that *f*′(*x*) = 0 is not a sufficient condition for *x* to be a minimum, because if *x* is a maximum then *f*′(*x*) = 0, and, if *f*″(*x*) = 0, then *x* can be an inflection point. Finding minima thus starts by calculating *f*′ and then solving *f*′(*x*) = 0.

In several dimensions, the “derivative” of a function *F*: ℝ^*D*^ → ℝ is the *gradient vector* 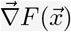, whose components are given by the partial derivatives of the function with respect to each component of 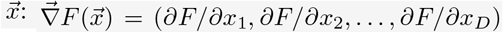. The gradient vector points in the direction of maximum variation of *F*, and is therefore zero (*i.e.*, equal to the null vector 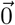) when 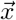 is a minimum:

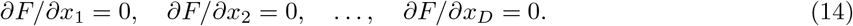

The derivative of a sum is the sum of the derivatives, so, deriving (13) with respect to, *e.g.*, *x*_*k*_, gives

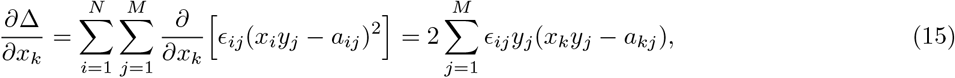

because *x*_*i*_*y*_*j*_ does not depend on *x*_*k*_ if *i* ≠ *k* and *a*_*ij*_ is constant for all *i* and *j*. Then,

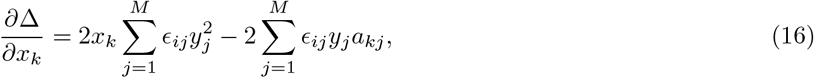

and the conditions *∂*∆/*∂x*_*i*_ = 0 and *∂*∆/*∂y*_*j*_ = 0 for all *i* and *j* yield

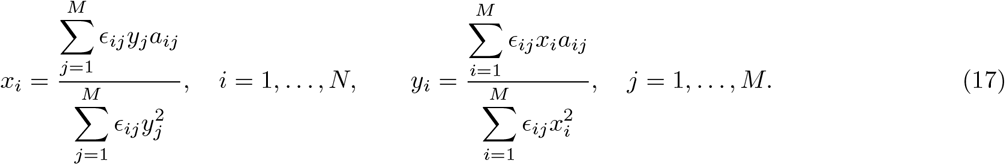

This is a system of *N* +*M* equations and unknowns that can be solved with different methods. Due to the large dimension of the systems arising in social interaction analysis, iterative method are often used. See Box 2.

### Box 2 Fixed point iterations

A point *x*\* ∈ ℝ is a fixed point of a function *f*: ℝ → ℝ if *f*(*x*\*) = *x*\*. Fixed points can be found under certain conditions^*a*^ by means of the iterative method

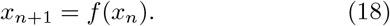

Starting from an initial point *x*_0_, the method builds a sequence *x*_1_, *x*_2_, *x*_3_, … that converges to (one of) the fixed point(s) of *f*; see Fig. 9. This method is also called method of successive approximations.

**Figure 9:**
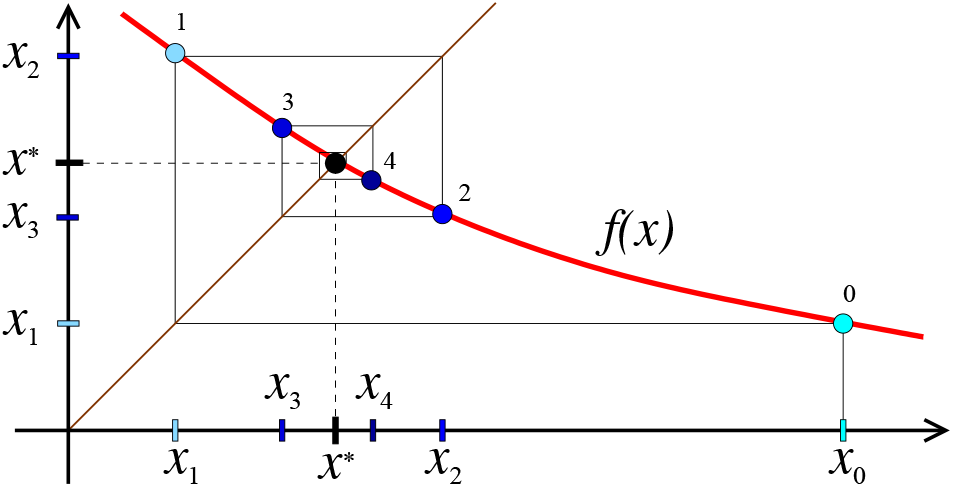
Iterations converging to (*x* x\**). Red line: function *f* (*x*); brown line: *y* = *x*; thin polygonal: iteration process; coloured dots: successive values of *x*_*n*_.

In *d* dimensions, a vector 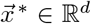 is a fixed point of a function 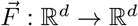 if 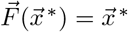.

When the dimension of the system *d* = *N* + *M* is very large, and in order to improve the stability of the recursion dynamics, it is convenient to use a relaxation method,

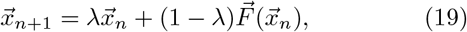

where *λ* ∈ (0, 1) is the weight of the previous iteration in the value of the new iteration, averaged with what would have been the new iteration.

## Footnotes

1 The average function of *a*(*x, y*) with respect to the variable *x* is given by 〈*a*〉_*x*_(*y*) = ∫_*x*_*p*(*x, y*)*a*(*x, y*)*dx*, where *p*(*x, y*) is the probability of occurrence of the state (*x, y*). However, force maps calculate the 〈*a*〉_*x*_ (*y*) as the mean of the values of *a*(*x, y*) over all the values of *x*, *i.e.*, 〈*a*〉_*x*_ (*y*) =∑_*x*_ *a*(*x, y*)/∑ _*x*_ 1, as if *p*(*x, y*) = 1 for all (*x, y*), because knowing *p*(*x, y*) is part of the problem.

a The function *f* must be contractive and the initial value of the iterations must be sufficiently close to the fixed point.

